# Elucidating the functional domain architecture of ArCS1, a biomineralizing myosin chitin synthase: I. The role of lipids

**DOI:** 10.64898/2026.08.31.748318

**Authors:** Sahar Nakhostin, Ingrid M. Weiss

**Author notes:** Correspondence: S.N., I.M.W.

## Abstract

In molluscs, chitin synthases are essential for biologically controlled biomineralization, with some variants possessing a myosin motor domain that may link polymer synthesis to the cytoskeleton.

Experimentally, we established a reliable workflow for expressing ArCS1_E22TM in *Dictyostelium discoideum* and developed effective purification methods to reconstitute ArCS1_E22TM in nanodiscs using MSPs and specific lipid composition. MSP1D1deltaH5 proved optimal for nanodisc formation, yielding homogeneous, monodisperse discs (∼8.2 nm). Lipids were refined to POPC:POPE:POPG (3:1:1) with 20% cholesterol, improving nanodisc quality and uniformity as observed by negative-stain EM. The full-length ArCS1 and its subdomains were modeled using AlphaFold3; the myosin motor, glycosyltransferase, and transmembrane regions are well-defined internally but loosely constrained relative to one another, suggesting flexible linking and conformational coupling. Modelling with Mg^2+^ and oleic acid as ligands and comparative analyses with bacterial cellulose synthase and yeast chitin synthase 1 provided insights into substrate binding and a potential mechanism for chitin polymerization and translocation.

This research establishes a standard procedure for comprehensive structural analyses of recombinant molluscan chitin synthase in near-native or biomimetic membranes. This sets the stage for high-resolution cryo-electron microscopy to determine the first experimentally resolved structure of a molluscan chitin synthase and to provide insight into the enzyme’s architecture and the regulatory mechanisms of biomineralization.

## Background

Molluscs with the ability to biomineralize their shells have evolved since 550 million years ago, combining an organic matrix secreted by the mantle that specifies where and how calcium carbonate phases nucleate, grow, and organize into hierarchical architectures such as nacre, prisms, and crossed lamellae (Knoll, 2003; Lowenstam and Weiner, 1989; Mann, 1983). Despite extensive study, the molecular mechanisms that link intracellular processes to extracellular matrix organization and mineral formation remain poorly understood. Within this matrix, chitin synthases generate the chitin scaffold that provides the primary polysaccharide framework and prefigures the microstructural order of the shell, thereby defining the architecture of the mineralizing interface (Marin et al., 2012; Weiss, 2012).

Comparative sequence and phylogenetic analyses indicate that molluscan chitin synthases share a conserved domain architecture that is likely essential for biomineralization-related chitin synthesis (Weiss, 2019; Zakrzewski et al., 2014). The catalytic apparatus with the classical GT-A fold belongs to the GT2 family of glycosyltransferases and contains several conserved motifs essential for catalysis, including aspartate residues in D/E, DxD, and GED motifs that coordinate nucleotide binding, cofactor binding, and acceptor deprotonation, respectively, as well as a Q/LxxRW pentapeptide, or the distinct NQRRRW variant in molluscan chitin synthases, that guides polymer transit through the catalytic channel (Bi et al., 2015; Gudnason et al., 2026; Nakhostin and Weiss, 2026). The myosin motor domain shows greatest similarity to class III myosins and includes a kinase-like N-terminus with motifs for ATP binding and catalysis; this domain generates directional mechanical force via actin-activated ATPase activity (Komaba et al., 2003; Weiss, 2012, 2019; Weiss et al., 2006). The chitin translocation domain comprises several transmembrane helices that form a channel extending from the catalytic center to the C-terminus (Weiss, 2019). Distinctive to molluscan CSs is a low-complexity and charge-rich domain at the C-terminal end that features a conserved SWGTRE motif (Weiss, 2019; Weiss et al., 2013). This region does not follow a typical globular domain structure; instead, it is predicted to be flexible and potentially form coiled-coil or self-associative regions that function as a sensitive “charge switch” (Weiss et al., 2013; Weiss and Marin, 2008). This specific switching point is physiologically relevant, as it closely corresponds to the natural pH range (7.2-7.5) of the mollusc’s extrapallial fluid, suggesting that the protein’s tendency to aggregate is fine-tuned to the environmental conditions of shell formation (Ghatak et al., 2013; Weiss et al., 2013).

Previous work has identified and analyzed the full-length chitin synthase sequence from mantle tissue of the genus *Atrina*, providing a molecular basis for further investigation (Weiss et al., 2006). However, understanding how these enzymes function requires structural insight into the molecular and mechanical cross-talk mechanism between material precursors and cells. Structural characterization of chitin synthase from *Atrina rigida* builds on established systems for recombinant expression in *Dictyostelium discoideum* and enables targeted analysis of its functional domains (Schönitzer et al., 2011; Weiss et al., 2013).

In this study, we aim to pave the way for 3D structure characterization of ArCS1 and its domains. First, we establish robust expression and purification of the extracellular domain ArCS1 (ArCS1_E22TM) in *Dictyostelium discoideum*, enabling biochemical and biophysical analysis. Second, we reconstitute purified ArCS1_E22TM into lipid nanodiscs assembled with membrane scaffolding proteins, creating a detergent-free, native-like membrane platform compatible with single-particle electron microscopy and sensitive to annular lipid interactions that often govern the stability and activity of processive synthases. Third, we leverage AlphaFold3 (Abramson et al., 2024) to model full-length ArCS1 and its subdomains, yielding testable hypotheses for domain organization, cytoskeletal coupling via the MMD, gating element positions, and polymer translocation geometry.

This integrated framework, expression, nanodisc reconstitution, and structure prediction, prioritizes the molluscan-specific questions that matter for shell biomineralization: How do the MMD and actin organize chitin trajectories at the membrane? Which extracellular segments of ArCS1 act as pH-tuned charge switches to bind mineral faces and prepattern nucleation? How do lipids shape the TM conduit to sustain processivity and regulate gating? By stabilizing ArCS1 in nanodiscs and mapping predicted interfaces, we set the stage for cryo-EM and functional assays to connect these features to chitin output and mineral templating.

## Results

### Expression and purification of recombinant ArCS1_E22TM

*Dictyostelium discoideum* Ax3-orf+ cell lines containing pDXA-E3TM-YFP plasmids were successfully expressing YFP-tagged ArCS1_E22TM, confirmed by confocal microscopy and YFP filter sets (Figure 1A). The Ax3-orf+ cell lines without plasmid transformation were used as a control for microscopy. For these samples, the basic settings for the detector gains of both fluorescence channels were applied on the confocal microscope, resulting in no fluorescence signal being detected. Cells were growing and behaving normally, moving around, and more than 90% of cells fluoresced. No specific change was observed in cells due to the recombinant expression of ArCS1_E22TM.

**Figure 1.**
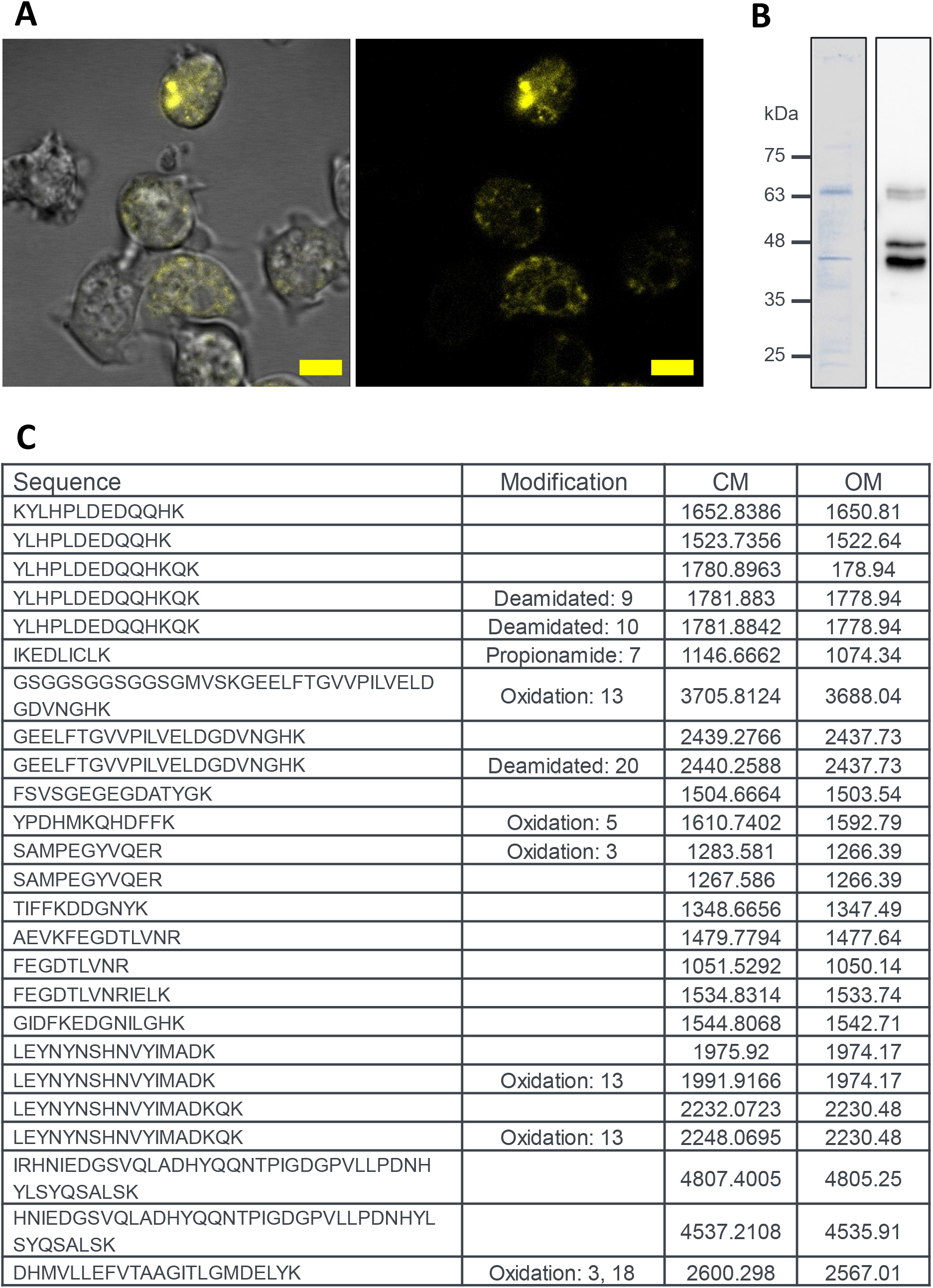
Expression and purification of the ArCS1_E22TM, (A) *Dictyostelium discoideum* expressing the extracellular domain of ArCS1, left image: overlay of YFP channel and DIC channels, right image: YFP channel. Scale bar, 5 µm. (B) Coomassie-stained gel and immunodetection of purified protein. The calculated molecular weight of ArCS1_E22TM is about 64 kDa, as detected in both analyses. However, a species at about 45 kDa was always detected for ArCS1_E22TM. (C) Identification of ArCS1_E22TM in detected bands in SDS-PAGE analysis of purified protein with Mass spectroscopy. Twenty-five different peptides obtained from an isolated ∼45 kDa protein band were identified and found to correlate with the sequence of ArCS1_E22TM, which has a theoretical molecular weight of ∼64 kDa. The calculated peptide mass (CM), observed peptide mass (OM), and peptide modifications are indicated: oxidation of methionine residues, deamidation of asparagine or glutamine, or propionamide modification of cysteine residues.

Different detergents and lysis conditions were tested to identify the optimal lysis condition and improve the stability of the protein of interest. The soluble recombinant homolog ArCS1_E22, derived from the extracellular domain of the mollusc myosin chitin synthase ArCS1, was previously shown to assemble into complexes at pH < 7.75 and could only be purified natively at pH 9 (Weiss et al., 2013). However, in a comparable experiment, lysis of cells with a pH 9 buffer led to the detection of truncated species, as had also been reported for ArCS1_E22. Similarly, the best result during lysis with detergents at neutral pH was obtained using 1% DDM. ArCS1_E22TM, a four-transmembrane-helix fragment with a calculated molecular weight of 63.8 kDa, displayed atypical SDS-PAGE behavior characteristic of a highly charged membrane protein; boiling induced aggregation, whereas heating at 55 °C enhanced detection without shifting the band position. Given that the high Asp/Glu content would be fully deprotonated at the elevated pH of the Tris-Glycine system, the intrinsic charge of the protein may interfere with migration depending on the extent of SDS binding, causing it to run at a molecular weight different from the predicted value (Marin and Luquet, 2007; Weiss et al., 2013). Switching to a Bis-Tris separation system and employing different staining methods did not alter the band position, but staining was markedly improved after eliminating background haze. Western blot analysis of the eluate (Figure 1B) supports this interpretation, revealing the characteristic bands of ArCS1_E22TM at approximately 45 kDa.

Ultimately, 2 mg of ArCS1_E22TM could be purified from 10 g of wet cell pellet, which is suitable for biochemical analysis and EM studies. To verify protein identity, individual bands from the immunoaffinity purification were subjected to in-gel tryptic digestion and mass spectrometric analysis. The characteristic band of ArCS1_E22TM at 45 kDa yielded identifications corresponding to peptides from the extracellular domain of chitin synthase with 40.5% coverage (Figure 1C). Coverage gaps correspond to predicted transmembrane helices and other hydrophobic regions, a well-known limitation of in-gel trypsin workflows when analyzing membrane proteins (Vit and Petrak, 2017). Minor oxidation and sporadic deamidation are common artifacts in this context and do not affect the overall identity conclusion. These routine technical modifications did not hinder confident sequence assignment. Under the instrument conditions employed, CM-OM differences generally fell within a few Daltons for larger peptides and approximately 1-2 Da for smaller peptides. However, no peptide corresponding to ArCS1_E22TM could be recognized for the 65 kDa band.

### ArCS1_E22TM nanodisc “Portraits”: assessing particle size and homogeneity

Based on the calculated surface area of 4 transmembrane helices of ArCS1_E22TM (560 Å^2^), MSP1D1deltaH5 with a nanodisc size of 8.2±0.6 nm could perfectly accommodate four TMHs, and there will be about 2.5 nm of lipids surrounding the protein of interest. In previous studies, a distance of 1-2 nm has been considered between the target protein and MSPs (Efremov et al., 2017; Li et al., 2024). Nanodiscs assembled from the mixed-lipid formulation with MSP1D1deltaH5 displayed a more homogeneous size distribution than those prepared with MSP1D1 (S01), indicating improved structural stability and uniformity of assembly, as confirmed by SDS-PAGE and size-exclusion chromatography analysis. Based on successful precedents, nanodiscs were assembled with a 3:1:1 molar ratio of POPC:POPE:POPG. This mixture’s phase behavior enabled reconstitution at 4 °C, thereby reducing protease activity and prolonging protein lifetime. Additionally, cholesterol, a major sterol in mantle lipids, was included in some experiments to modulate membrane fluidity and improve protein folding, while bare nanodiscs served as controls.

Comparing the bare nanodiscs and ArCS1_E22TM: MSP1D1deltaH5:PL chromatogram (Figure 2A), the elution assemblies take place at almost identical column volume, with ArCS1_E22TM:MSP1D1deltaH5:PL having a shoulder at the left side of the peak, whereas empty NDs depict a symmetrical peak. This indicates either a minor higher-molecular-weight population, such as partially aggregated species, or incomplete resolution between different species. The latter occurs when the main population and the larger species are close in size; the column may not fully resolve them, producing a broad-front or shoulder peak rather than two distinct peaks. Usually, if the membrane protein reconstituted in NDs does not have a big extra- or intracellular domain and the main domain is inside the lipid membrane, the empty ND and the reconstituted ones elute at the same column volume (Vilela et al., 2024). This is true for ArCS1_E22TM, which has four TMHs embedded in the membrane, a low-complexity region at the extracellular domain, and only a YFP domain intracellularly. Although employing SEC is a common practice to separate empty NDs from full ones before EM, having empty NDs in EM micrographs will not interfere with subsequent structural data analysis.

**Figure 2.**
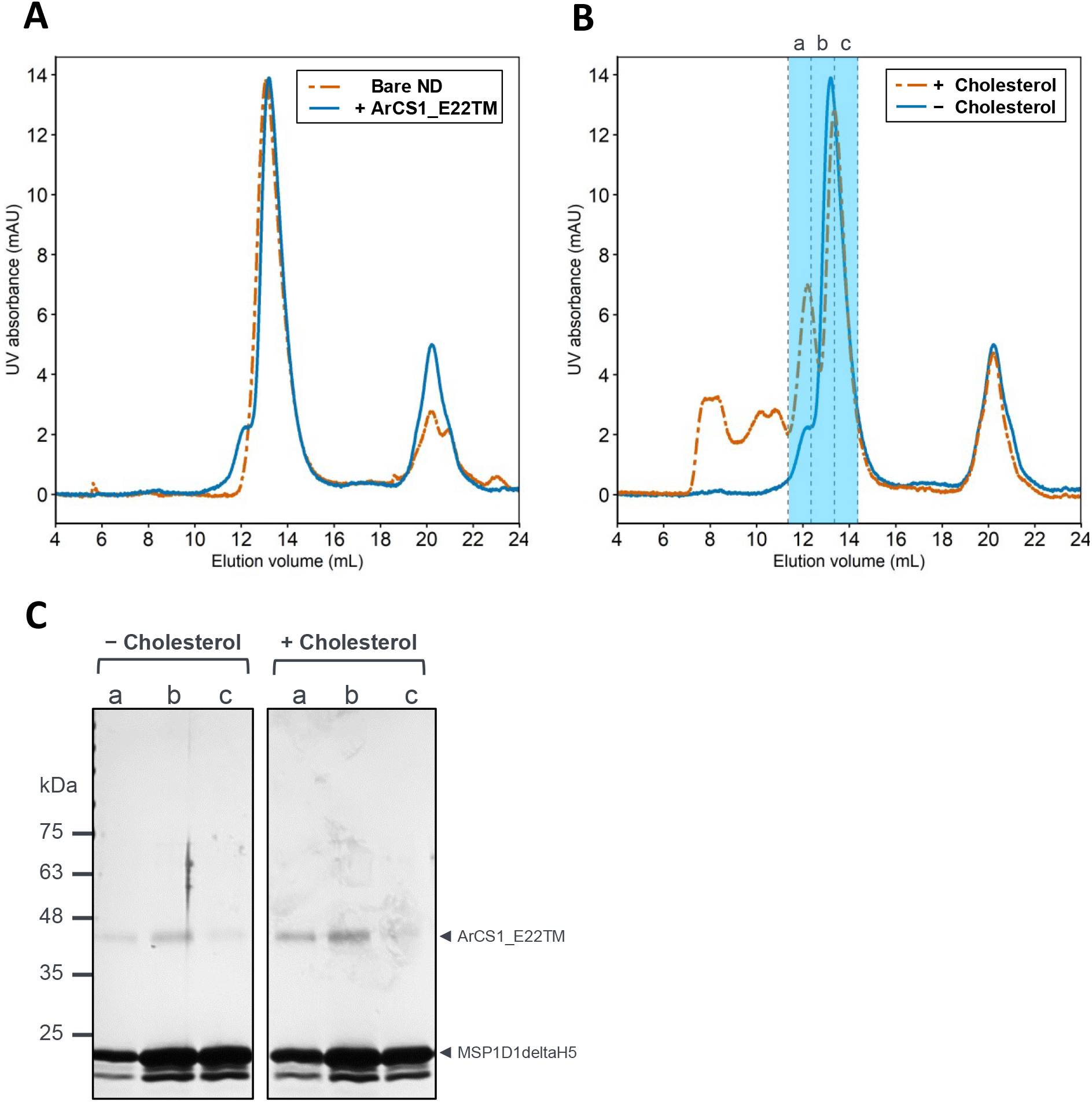
Reconstitution of ArCS1_E22TM in nanodiscs formed with MSP1D1deltaH5 and mixed-lipid formulation, (A) SEC chromatogram of bare ND and ArCS1_E22TM:MSP1D1deltaH5:PL, (B) SEC chromatogram of ArCS1_E22TM:MSP1D1deltaH5:PL with/without cholesterol, (C) SDS-PAGE analysis of ArCS1_E22TM:MSP1D1deltaH5:PL with/without cholesterol, fractions are marked in (B).

Comparative analysis of nanodiscs prepared in the presence and absence of cholesterol provided further insight into its potential contribution to nanodisc stability and the conformational state of the incorporated membrane protein. Under identical conditions of reconstituting NDs with a molar ratio of 5:20:980, ArCS1_E22TM:MSP1D1deltaH5:Lipids, the elution profile of NDs containing cholesterol showed distinctive characteristics (Figure 2B). There are 2-3 peaks at higher molecular weights, which can be assigned to the cholesterol-containing liposomes for several reasons. Initially, these peaks were consistently observed in all SEC elution profiles of NDs containing cholesterol. Furthermore, no protein was stained for the fractions corresponding to these peaks by SDS-PAGE (Figure 2C). On the other hand, although the cholesterol’s main UV absorbance happens to be in wavelengths below 210 nm, there have been reports indicating weak peaks at 283 nm (Heilbron et al., 1927). Additionally, given the significantly higher lipid concentration relative to protein in the reconstituted mixture, the earlier peaks in the chromatogram of ArCS1_E22TM:MSP1D1deltaH5:PL with cholesterol were attributed to cholesterol-containing liposomes.

What is worth noting is that what was earlier regarded as the left shoulder of the main peak in the ArCS1_E22TM:MSP1D1deltaH5:PL chromatogram has been replaced by a more pronounced peak in the nanodiscs with cholesterol chromatogram. Moreover, the stained bands for ArCS1_E22TM in SDS-PAGE analysis of fractions are faint for fraction a in the complexes without cholesterol and more distinct in complexes with cholesterol. This agrees with SEC results, which show a second peak for ArCS1_E22TM:MSP1D1deltaH5:PL assemblies with cholesterol, suggesting the presence of two distinct species of different sizes.

Negative-stained micrograph of both complexes (Figure 3A-B) depicts homogenous and mainly monodisperse particles for NDs with/without 20% cholesterol. Particle size was measured for fraction B of both assemblies, and the data reveal a primary population of particles with a Gaussian distribution (Figure 3C-D). While a minor fraction of smaller particulates (approx. 5-7 nm) was observed for ArCS1_E22TM:MSP1D1deltaH5:PL assemblies, the dominant population is highly uniform. After excluding outliers (more than 2 standard deviations), the core population was calculated to be 9.47±0.94 nm. Assemblies with cholesterol were more homogenous with an average size of 8.2±0.88 nm, indicating a smaller size range and falling perfectly within the MSP1D1deltaH5 ND size range as reported before (Hagn et al., 2013). By taking a closer look at the size distribution of particles and their elution profile, one can easily note that the elution of both groups of particles happens almost at the exact column volume. Still, there is a 1-2 nm difference in the average particle size, proving that the SEC column cannot separate particles with this slight size difference.

**Figure 3.**
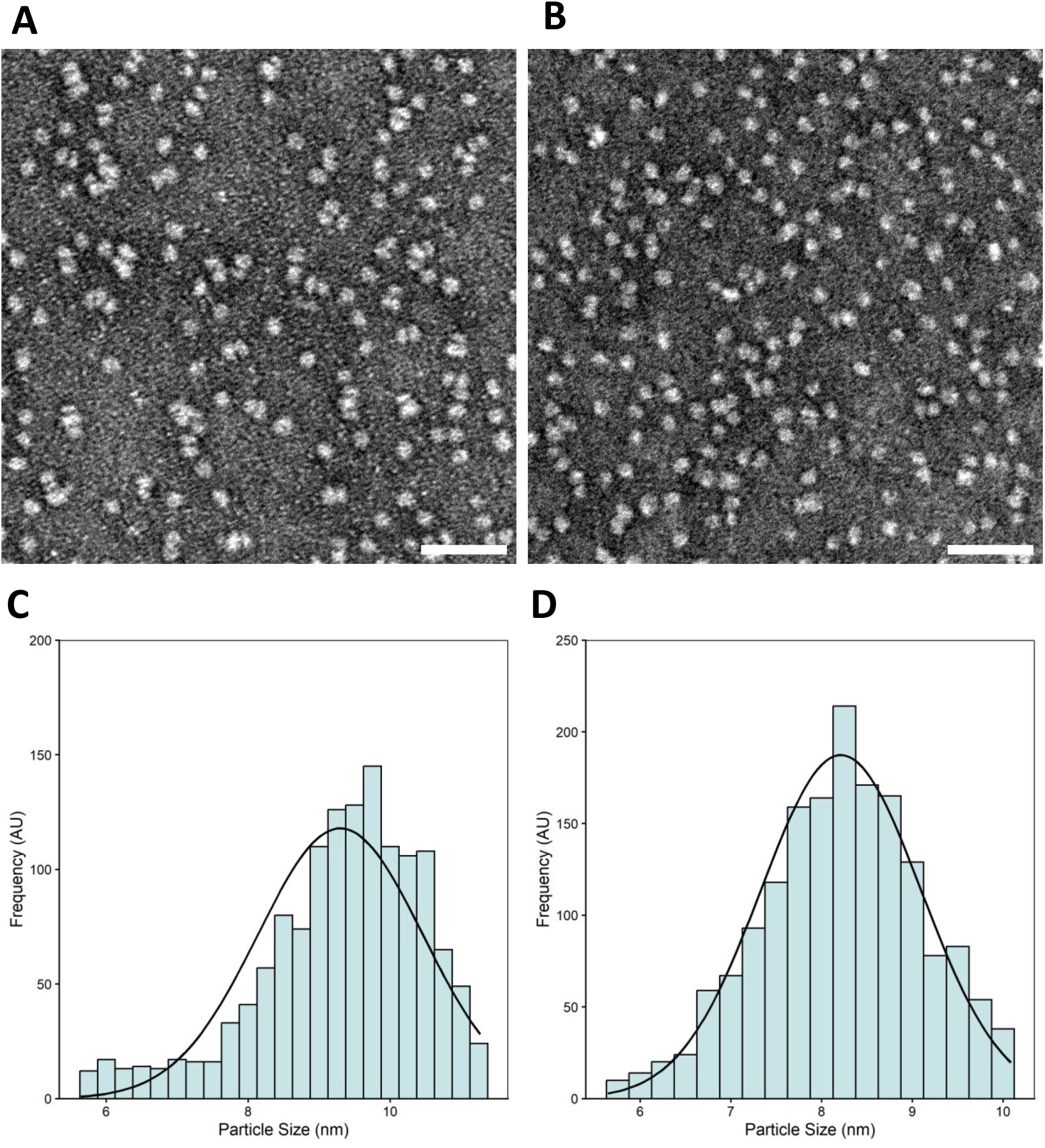
Morphological and size distribution analysis of MSP1D1deltaH5 ND with mixed-lipid formulation, (A) Representative negative-stained micrograph of ArCS1_E22TM:MSP1D1deltaH5:PL, (B) Representative negative-stained micrograph of ArCS1_E22TM:MSP1D1deltaH5:PL with 20% cholesterol. Micrograph scale bars, 50 nm. (C) Particle size distribution histogram ArCS1_E22TM:MSP1D1deltaH5:PL, (D) Particle size distribution histogram ArCS1_E22TM:MSP1D1deltaH5:PL with 20% cholesterol.

The addition of 20% cholesterol appears to decrease the average nanodisc size and increase their homogeneity. This could stem from cholesterol’s influence on lipid packing within the nanodisc or its effect on the interaction between the protein and the lipid bilayer. The monodisperse nature of nanodisc preparations is crucial for downstream applications, enabling more accurate and reliable biophysical characterization and functional assays (Efremov et al., 2017; Serna, 2019). The SEC results suggest that the size difference between nanodiscs with and without cholesterol is subtle. However, the presence of a minor fraction of smaller particulates in the sample without cholesterol might indicate some instability or dissociation of the complex. The improved homogeneity of cholesterol-containing nanodiscs is a critical factor for high-resolution structural studies. Overall, NDs made with MSP1D1deltaH5 and lipid combination of POPC, POPE, POPG, and 20% cholesterol appear promising for continuing the research.

### ArCS1 structure in silico

AlphaFold3 predicted the structure of chitin synthase from *Atrina rigida*, annotating a myosin motor, GT domain, 18 TMHs, and disordered regions. The model shows a membrane protein topology with TMH clusters, a cytosolic GT domain, and a cytosolic myosin motor domain, with Mg^2+^ included as a ligand (S02). Confidence (pLDDT) is high for MMD, TMHs, and the GT domain, but low for low-complexity and extracellular regions. The PAE heatmap reveals a modular organization in which individual domains are well-defined but weakly constrained relative to one another. To mimic the membrane environment, oleic acid molecules were added in AlphaFold3 runs, resembling unsaturated fatty acids in marine bivalve membranes (Toghani et al., 2025). The resulting PAE map still showed well-folded domains with flexible linkers, but adding 30 oleic acids improved the coherence of the transmembrane region and raised the pTM score from 0.51 to 0.58. This moderate increase suggests better constraint on membrane-embedded parts, though uncertainty remains for such large, multi-domain proteins with disordered segments.

To contextualize the myosin motor domain of the model, the predicted myosin domain from ArCS1 was superposed onto the crystallographic model of human myosin 1c (Figure 4A). It should be noted that the MMD of ArCS1 exhibits the highest similarity to the Myo III family. However, no 3D structure is available for any Myo III, so human myosin 1c (Myo1c, PDB entry: 4BYF) was chosen instead. Myo1c is a suitable candidate for structure superposition and 3D structural alignment with the MMD of ArCS1. On one hand, it shares a high degree of homology and structural similarity with ArCS1_MMD, making it a relevant template for comparison. On the other hand, Myo1c and ArCS1_MMD may exhibit functional parallels, as class I myosins, including Myo1c, promote the formation and elongation of actin protrusions. For example, Myo1c has been observed to localize at the tips of actin structures and is essential for the mechanical processes that enable membrane movement and protrusion dynamics, thereby facilitating the formation and elongation of these structures (Komaba and Coluccio, 2010). Additionally, class I myosins contribute to processes such as membrane tension regulation, further supporting the development of actin-driven protrusions (Houdusse and Titus, 2021). Core secondary-structure elements, including several helices and beta strands, align closely and superpose with near-atomic agreement, indicating a conserved architectural scaffold. In other segments, corresponding residues deviate by approximately 1-10 Å, consistent with local insertions, surface loop differences, or hinge-like flexibility between subdomains. These offsets are enriched in regions that show moderate pLDDT and elevated PAE in the predicted model, suggesting that the observed displacements likely reflect genuine conformational variability rather than global misfolding. The overall correspondence supports a shared myosin fold while highlighting species-specific adaptations that may modulate the coupling between nucleotide binding, actin interaction, and membrane-associated functions in ArCS1.

**Figure 4.**
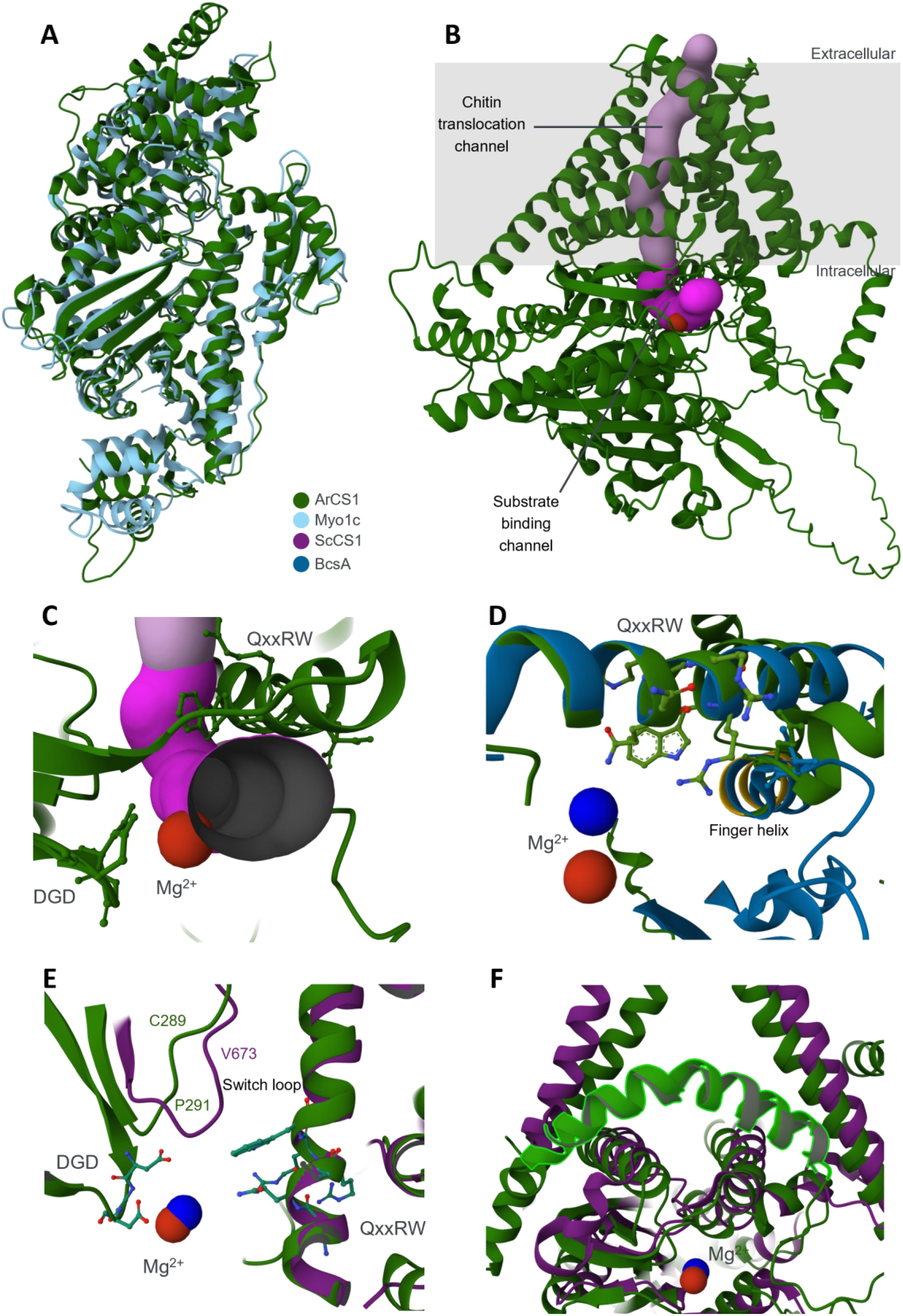
Structure assessment of full-length ArCS1 predicted by AlphaFold3. (A) Superposed structures of the predicted MMD from ArCS1 and human Myo1c. The core helices and beta strands superpose closely, while localized differences of approximately 1–10 Å highlight insertions, surface loop variability, and potential hinge-like motions between subdomains. (B) ArCS1 model displaying two critical channels (C) The substrate binding channel (magenta) for UDP-N-acetyl-α-D-glucosamine entry, at the proximity of QRRRW and DGD motifs. (D) Structure alignment of ArCS1 and BcsA GTD, a close look at “finger helix” (yellow) with UDP-GlcNAc bound to BcsA. (E) Structure alignment of ArCS1 and ScCS1, comparison of the possible switch loop motif. (F) Highlighting the curved helix directing the ECD to the cytosolic space. In all images, Mg^2+^ of ArCS1 is colored red. The QxxRW and DGD motifs are represented as ball-and-stick models.

Examining the GT domain more closely (Figure 4B-C), the nucleotide-binding domain containing the DGD motif can be identified near Mg²⁺, which most likely mediates the binding of Mg²⁺ to the aspartic acid carboxylates, which then form complexes with the phosphate oxygens of UDP-activated sugar substrates. In addition to the DGD motif, the polymer-guiding/channeling motif “QRRRW” is found near Mg²⁺, within a formed channel that may serve as the chitin synthesis and transport channel.

Based on the AlphaFold3-predicted structure, two channels were identified (Figure 4B), revealing the structural features of chitin synthase. The substrate-binding channel, colored magenta in Figure 4, serves as the entry point for the nucleotide sugar substrate (UDP-N-acetyl-α-D-glucosamine). This channel begins on the cytosolic side and is lined by the DGD and QRRRW motifs, enabling interactions with the hydroxyl groups of the sugar moiety and assisting in its proper orientation for catalysis. It then connects to a pocket located between the helices arranged in the transmembrane region, extending into the extracellular space. The translocating channel may contain flexible regions or loops that act as “gates” to control the passage of the polymer chain, similar to mechanisms reported in other CSs (Chen et al., 2023, 2022). These regions can undergo conformational changes to prevent premature release of the chitin strand, effectively guiding it toward its final destination.

The substrate-binding channel contains significant hydrophilic regions, with a diameter of approximately 4-6 Å, optimized to accommodate the polar substrate UDP-N-acetyl-α-D-glucosamine and facilitating essential interactions for substrate stabilization. This channel extends about 35 Å in length, providing a sufficient pathway for substrate transit. In contrast, the chitin-translocating channel exhibits a mix of hydrophobic and hydrophilic features, with a variable diameter of 4-7 Å and a length of approximately 54 Å, allowing it to engage with the growing hydrophobic chitin polymer while also accommodating water, ensuring efficient polymer extrusion.

To explore the detailed features of the domains, the model was aligned with bacterial cellulose synthase (BcsA, PDB entry: 5EIY) and *S. cerevisiae* CS1 (ScCS1, PDB entry: 8K3V) in complex with UDP-GlcNAc. In the alignment of ArCS1 and BcsA (Figure 4D), part of the core beta sheets show very close positioning. However, helices in the mollusc CS align only moderately with those in cellulose synthase, indicating several major differences between cellulose- and chitin-synthesizing systems. Meanwhile, it is worth noting that the suggested concept of a “finger helix,” presented by Jochen Zimmer’s team (Morgan et al., 2016), may also apply to ArCS1. Based on their experimental evidence, a finger helix stabilizes the chain during the pre-elongation phase. This finger adjusts its position after chain elongation (pre-translocation) and then “snaps” back to its original position while remaining attached to the newly added sugar. This retraction signifies the new non-reducing acceptor state and prepares the enzyme for the next cycle (post-translocation). By comparing the ArCS1 model and the structure of BcsA, a similar helix with very close alignment to the finger helix of BcsA near the substrate-binding pocket is found (Figure 4D). Isoleucine “I340” identifies the finger helix in BcsA. In ArCS1, there is also an isoleucine at position 1642, followed by a highly conserved motif featuring the “ED” doublet, which is common to both BcsA and mollusc chitin synthases (Weiss, 2019). Considering that a similar finger helix can be recognized for ArCS1, the switchable nature of D-terminated domains in response to pH changes is particularly interesting for biomineralization processes, where ion gradients across membranes may significantly influence control mechanisms at the mineralizing interface.

In the alignment between the modeled ArCS1 and ScCS1, both proteins show good alignment in many of their alpha helices and beta sheets, particularly in the core of the GT, indicating comparable folding patterns and functional domains. For ScCS1, a switch loop has been reported that plays a pivotal role in regulating access to the active site of the CS. The loop, particularly the “VLPGA” motif, undergoes a significant conformational change upon substrate binding. When UDP-GlcNAc and GlcNAc bind, the switch loop shifts away from the active site, allowing a continuous path for chitin elongation and facilitating the transfer of the growing polymer through the channel (Chen et al., 2023). By analyzing the sequence and structure alignment of both proteins, a very similar loop can be confidently deduced for the model of ArCS1 (Figure 4E). In ArCS1, the “CCPGC” motif aligns very well with the “VLPGA” motif of ScCS1 and shows an analogous structure that could serve as a comparable mechanism to the switch loop of ScCS1. Nevertheless, all these possible mechanisms can only be proven by further in-depth studies of ArCS1 in different conformational stages and through mutagenesis.

In the sequence alignment of ArCS1 and ScCS1, the region responsible for helix 13 in ArCS1 displays high similarity to ScCS1 helix 5, also showing high similarity in structure superposition, both being curved in the membrane region (Figure 4F). For ScCS1, this curved helix continues intracellularly, comprising a segment of approximately 56 residues, mostly disordered but containing a “swapping loop” (Chen et al., 2023). This loop plays a critical role in ScCS1 dimerization and contributes to the interface between the two protomers of the enzyme.

While it is possible to infer an extracellular domain for molluscan chitin synthase using several standard bioinformatic tools and UniProt annotations, AlphaFold suggests that helix 13 curves inward, positioning the extracellular domain in the cytosol. This domain, rich in highly charged amino acids and characterized as a low-complexity region, poses a challenge for AlphaFold, which produces a structure with very low confidence.

AlphaFold may generate misleading structural order in disordered regions (Abramson et al., 2024). These areas are usually indicated as having very low confidence. Additionally, the performance of AlphaFold is heavily dependent on sequence similarity; proteins with little or no homology to characterized structures may yield unreliable predictions (Jones and Thornton, 2022). Consequently, AlphaFold may not fully capture the conformational diversity required to understand the complexities of unique proteins. Moreover, current assessments indicate that AlphaFold has not been comprehensively validated for predicting the effects of mutations on protein structure or for providing accurate protein models in specific cellular contexts, such as varying pH, as is the case in biomineralization. On the other hand, membranes display a certain degree of curvature (Gamage and Pan, 2025; Vanni et al., 2014), especially at the tips of microvilli, where chitin synthase is likely to be active (Weiss, 2019). Therefore, it is crucial to recognize that transmembrane regions are actively shaped, and standard predictions of membrane topology may not capture the full picture.

## Discussion

This work establishes a reproducible workflow for expressing, purifying, and reconstituting the extracellular domain of molluscan chitin synthase ArCS1_E22TM from *Atrina rigida* into nanodiscs, enabling downstream cryo-EM studies. The purification of ArCS1_E22TM posed several challenges due to its unique biochemical properties. To address these, a purification strategy employing immunoaffinity purification followed by Pure GFP elution, along with careful buffer optimization, was adopted to achieve high selectivity and reduce background contamination. The protein’s high density of charged amino acid residues interferes with SDS binding and electrophoretic mobility, contributing to its unusual behavior. Consistent with this, studies suggest the protein species exhibit a low affinity for SDS, aligning with their low hydrophobic amino acid content (Weiss et al., 2013). It is speculated that the N-terminus of the soluble homolog of ArCS1_E22TM, containing the SWGTRE motif and coiled-coil region, controls whether ArCS1_E22 is more likely to be found as associated multimers, individual monomers, or in complexes with host proteins(Weiss et al., 2013).

The successful reconstitution of ArCS1_E22TM into nanodiscs required careful optimization of several parameters, including the choice of MSP scaffold, lipid composition, and reconstitution conditions. Initial attempts yielded heterogeneous nanodisc populations, underscoring the importance of tailoring these parameters to the specific properties of the target protein. MSP1D1deltaH5 could better accommodate the four transmembrane helices of ArCS1_E22TM while providing sufficient space for lipid molecules to shield the protein from the MSP.

The lipid composition of the nanodiscs was carefully chosen to mimic the natural environment of ArCS1_E22TM. The cell membranes of marine molluscs, particularly mussels, are rich in phospholipids, with POPC and POPE being major components of mantle tissues (De Moreno et al., 1980; Misra et al., 2002). POPC offers bilayer stability and neutral charge, while POPE introduces curvature stress that is manageable via lipid-to-MSP ratios. POPG contributes a negative surface charge that influences protein-lipid interactions and can modulate membrane protein oligomerization states (Dowhan, 1997; van Meer et al., 2008). Moreover, several successful nanodisc preparations with a 3:1:1 POPC:POPE:POPG ratio have been reported in previous studies (D. Goddard et al., 2015; Nadezhdin et al., 2023; Shen et al., 2023, 2022).

Given that sterols, with cholesterol as the major component (up to 30% in some species), are a consistent and significant part of mantle lipid composition in marine molluscs (De Moreno et al., 1980), cholesterol was included in the nanodisc assembly experiments. This decision was based on the understanding that cholesterol can modulate membrane fluidity and protein function (Gu et al., 2019; Nes, 1974; Regen, 2022), potentially improving the folding of the target protein ArCS1_E22TM. The inclusion of cholesterol significantly impacted the resulting nanodiscs, as cholesterol-containing nanodiscs were smaller and more homogeneous than those without cholesterol, suggesting that cholesterol promotes tighter lipid packing and may influence the interaction between ArCS1_E22TM and the lipid bilayer. Overall, the results indicate that the optimized nanodisc assembly protocol, using MSP1D1deltaH5 in combination with a mixed lipid formulation of POPC, POPE, POPG, and 20% cholesterol, yields homogenous and stable nanodiscs suitable for further structural and functional characterization of ArCS1_E22TM.

Parallel AlphaFold3 modeling predicted a multi-domain architecture comprising an intracellular myosin motor domain, a cytosolic glycosyltransferase catalytic core, and transmembrane helices forming a translocation channel. Using oleic acid to mimic the membrane environment during protein modeling enhances the confidence of transmembrane region predictions, leading to better domain packing. Although oleic acid is not a perfect match for molluscan cell membranes, this method demonstrates that incorporating lipid context into structure prediction can improve residue and interface prioritization for experimental validation, especially at GT-TM interfaces, where catalysis and chitin translocation are likely regulated. The MMD of ArCS1 shares structural similarities with class I myosins, suggesting a role in interacting with cytoskeletal elements to influence shell formation. This supports a model where motor-domain dynamics spatially regulate chitin deposition, potentially guiding it along the shell seam or at repair fronts. The extracellular domain, predicted by AlphaFold3 to be located within the cytosolic space and exhibiting low pLDDT scores, is a region of interest due to its potential regulatory role influenced by pH changes, mineral ions, and shell matrix components. Its charged sequences suggest involvement in biomineralization, protein-mineral interactions, and matrix organization, possibly acting as a scaffold at the mineralization front rather than having direct catalytic function.

The integration of experimental and computational data supports a model in which chitin polymerization by the GT domain is coupled to extrusion through a membrane-embedded channel, regulated by gating loops and modulated by the MMD for cytoskeletal coordination, with the extracellular domain poised to respond to biomineralization cues. However, further high-resolution cryo-EM and functional studies across conformational states are needed to validate these mechanistic proposals.

## Conclusion

This work advances our understanding of molluscan myosin-chitin synthases by integrating computational predictions with membrane-aware biochemical and biophysical analyses. The convergence of domain architecture with functional hypotheses highlights ArCS1 as a compelling model for studying how enzymatic catalysis, mechanical force, and membrane topology converge to orchestrate biomineralization. The study lays a solid groundwork for future high-resolution structural work and functional dissection, and it contributes a generalizable framework for the integrative analysis of complex membrane enzymes central to biomineralization processes.

For a biology audience, the message is straightforward: molluscan chitin synthases are not generic carbohydrate polymerases. Their myosin fusion, conserved catalytic pore, lipid-sensitive translocation pathway, and pH-responsive extracellular modules together form a shell-building machine that links intracellular force generation to extracellular mineral choreography. The approach presented here provides immediate experimental access to that machine and a roadmap for testing how ArCS1’s domains cooperate to control the timing, orientation, and placement of chitin at the mineralizing front.

## Materials and methods

All procedures were performed according to established protocols, those provided by the Dicty-Base web resource (www.dictybase.org), or the manufacturer’s recommendations.

### Cell cultivation and ArCS1_E22TM screening

Spores of well-expressing transformants of ArCS1_E22TM were readily available from the previous research (Weiss et al., 2013). Briefly, *Dictyostelium discoideum* cells were cultured in HL5 media supplemented with 30 µg/mL G418 (Roth) either in shaking cultures at a density of 1×10^4^ - 4×10^6^ cells/mL or in petri dishes, keeping the density of the cells between 1×10^4^ and 2×10^7^ cells/mL at 22 °C. The fluorescence signals of YFP-tagged ArCS1_E22TM were detected in an LSM 710 Meta confocal microscope (Zeiss) while cells were in LoFlo medium. YFP was excited by a 514 nm laser and emitted a fluorescence signal at 527 nm. Imaging parameters were adjusted to maximize spatial resolution based on fluorescence intensities significantly higher than those observed in non-transformed cells. Image processing was carried out using the Zeiss Zen software package. In the confocal datasets, signals are represented in yellow or grayscale, corresponding to photomultiplier tube signals originating from YFP emission.

### ArCS1_E22TM purification and detection

Cells were harvested by centrifugation at 1000 ×g and 4 °C (Rotanta 460R, Hettich) and washed twice with 1× PBS. About 4 g of cells were resuspended in 8 mL of lysis buffer (100 mM Tris-Cl pH 7.4, 150 mM NaCl, 40 mM MgCl_2_, 1% (w/v) DDM), snap-frozen in liquid nitrogen, and gently thawed on ice. Then they were homogenized using a dounce-homogenizer for 5-6 times. To remove cell debris, they were centrifuged at 1000 ×g for 10 minutes at 4 °C. The supernatant was then subjected to magnetic beads (Dynabeads, Thermo Fisher) bound to an anti-GFP antibody from mouse IgG1 (Roche). The complex was rotated gently overnight at 4 °C. The next day, purification was continued by collecting the supernatant and washing the magnetic bead twice with the wash buffer (20 mM Tris-Cl pH 7.4, 150 mM NaCl, 8 mM MgCl_2_, 0.2% (w/v) DDM). Then, the antigen-antibody complex was eluted with GFP elution buffer (20 mM Tris-Cl pH 7.4, 150 mM NaCl, 0.2% (w/v) DDM, 0.25 mg/mL GFP) and rotated overnight at 4 °C. Subsequently, the eluate was separated from the beads using a magnet, filtered through a 0.22 μm syringe filter, and subjected to size-exclusion chromatography on a Superdex Increase 10/300 GL column (Cytiva) at a flow rate of 0.35 mL/min.

All fractions were analyzed by SDS-polyacrylamide gel electrophoresis (SDS-PAGE) and Western blotting. An equal amount of fractions was heated in Laemmli buffer for 30 min at 55 °C prior to electrophoretic separation using 10% gels. Coomassie Brilliant Blue R-250 or silver staining, with detection limits of 50 ng and 1 ng per band, respectively, was used for detection. For western blotting, electrophoretically separated proteins on the gel were equilibrated in transfer buffer (6.06 g/L Tris, 3.09 g/L Boric acid, 10% (v/v) Methanol) for 15 min; the blotting sandwich was assembled and run for 75 minutes at 20 V and 300 mA. After blocking, membranes were incubated with primary antibody (Anti-GFP IgG1, Roche) and with the secondary antibody (Anti-Mouse, Rockland), respectively. After washing, detection was carried out with the Luminata Crescendo Western HRP substrate (Merck) and imaged using a Fusion Edge X7 Imaging System (Vilber Lourmat).

### Peptide fingerprinting and mass spectrometry

The bands of interest were excised, cut into ∼1 mm³ cubes, and transferred to microcentrifuge tubes. Gel pieces were destained with 100 µL of 100 mM ammonium bicarbonate/acetonitrile (1:1) for 30 minutes with occasional vortexing, then shrunk with 500 µL of acetonitrile until white. After removing acetonitrile, enough trypsin buffer (13 ng/µL in 10 mM ammonium bicarbonate with 10% acetonitrile) was added to cover the dry pieces. After 30 minutes, additional buffer was added if needed; the pieces were saturated for 90 minutes, then covered with 10-20 µL ammonium bicarbonate buffer and incubated overnight at 37 °C. Following digestion, 100 µL of extraction buffer (1:2, 5% formic acid/acetonitrile) was added (adjusted to the gel volume) and incubated for 15 minutes at 37 °C with shaking. The supernatant was collected, dried in a vacuum centrifuge, and sent to the mass spectrometry facility. Spectra were searched against a project-specific FASTA tailored to *Dictyostelium* and ArCS1 constructs.

### Reconstitution of ArCS1_E22TM in Nanodiscs

MSP1D1deltaH5 proteins were expressed and purified as described in previous studies (Hagn et al., 2013). Purified MSP1D1deltaH5 was concentrated using a 10,000 MWCO filter concentrator, with final protein concentration determined by A280 measurement (ε=21,430 M^-1^⋅cm^-1^).

Stock solution of lipids POPC, POPE, POPG, and Cholesterol (Avanti Polar Lipids) was prepared in chloroform, and the phosphate quantification assay was performed to determine the exact concentration of lipids (Fiske and Subbarow, 1925). The chloroform was evaporated under the gentle stream of nitrogen gas, and desiccated overnight under vacuum. The lipid mixture was rehydrated in 20 mM Tris-HCl, pH 7.4, 100 mM NaCl, 0.5 mM EDTA, and 50 mM sodium cholate. For reconstitution into nanodiscs, purified ArCS1-E22TM thereof in DDM micelles (Cube-biotech) were combined with hydrated lipid and MSP at the following molar ratios: ArCS1_E22TM:lipid:MSP, 5:20:980 with/without 20% cholesterol. The mixtures were incubated at 4 °C for 1 hour. Detergent removal was facilitated by the addition of 0.8 g/mL Biobeads (Amberlite™ XAD®-2, Sigma) followed by incubation at 4 °C overnight. Biobeads were then removed using a 0.45 µm filter. Size exclusion chromatography was employed to segregate nanodiscs with ArCS1-E22TM from empty nanodiscs in disc-forming buffer (20mM Tris-HCl, pH 7.4, 100 mM NaCl, 0.5 mM EDTA). Fractions were concentrated using a 100,000 MWCO filter concentrator (Amicon, Merck) and assessed via SDS-PAGE to verify reconstitution and pull out ArCS1-E22TM-containing nanodiscs.

### Negative staining electron microscopy

Copper grids (Mesh 300) were washed in HCl and EtOH, coated with a Formvar layer, and then coated with thin (50-100 Å) amorphous carbon films. Grids were glow-discharged, and 5 μL of the sample was deposited on a grid for 30 seconds. Then, the grid was washed twice with 100 μL of disc-forming buffer instantly. The excess water was blotted with filter paper, and the specimen was stained three times with 10 μL of 2% uranyl acetate. Samples were imaged at room temperature using a Zeiss EM10 electron microscope (Zeiss) equipped with a LaB_6_ filament and operated at an acceleration voltage of 60 kV. Particle sizes were measured with ImageJ (Schneider et al., 2012). Statistical analysis was conducted in R 4.5.2 (R Core Team, https://www.R-project.org/). An iterative Gaussian fit determined the main particle population’s size and uniformity; particles beyond two standard deviations from the raw mean were excluded from final mean and SD calculations.

### Structure prediction of ArCS1

AlphaFold3 (https://alphafoldserver.com/) was utilized to predict the three-dimensional structure of the full-length ArCS1 protein and its sub-domains. The amino acid sequence was obtained from the UniProt database (The UniProt Consortium, 2025) and subsequently input into AlphaFold, with/without ligands. The quality of these models was assessed through the confidence scores provided by AlphaFold. Visualization of the predicted structures was conducted using Mol*Viewer (Sehnal et al., 2021). Comparative analyses were performed with existing experimental structures sourced from the Protein Data Bank (Berman et al., 2000) to identify structural similarities and differences. Bioinformatics tools, such as MoleOnline (Raček et al., 2025) and UCSF ChimeraX (Meng et al., 2023), were employed to further analyze the structural features and explore molecular interactions.

## Abbreviations

CS: Chitin synthase
GlcNAc: N-Acetylglucosamine
UDP: Uridine diphosphate
TMH: Transmembrane helix
TMD: Transmembrane domain
GT: Glycosyltransferase
ECD: Extracellular domain
MMD: Myosin motor domain
ATP: Adenosine triphosphate
TEM: Transmission electron microscopy
CMC: Critical micelle concentration
MSP: Membrane scaffolding protein
SEC: Size exclusion chromatography
G/YFP: Green/Yellow fluorescent protein
PL: Phospholipid
ND: Nanodisc
pLDDT: Predicted local distance difference test
pTM: Predicted template modeling
PAE: Predicted aligned error

## Acknowledgment

Generous financial support by the Carl Zeiss Foundation for the infrastructure project ChitinFluid (project no. P2019-02-004) was gratefully acknowledged. We thank the mass spectrometry group at the Regensburg Center of Biochemistry (RCB, University of Regensburg) for protein mass spectrometric analyses.

