## Supplementary material for "Elucidating the functional domain architecture of ArCS1, a biomineralizing myosin chitin synthase: I. The role of lipids": Suplementary information 1&2

### Supplementary information 01

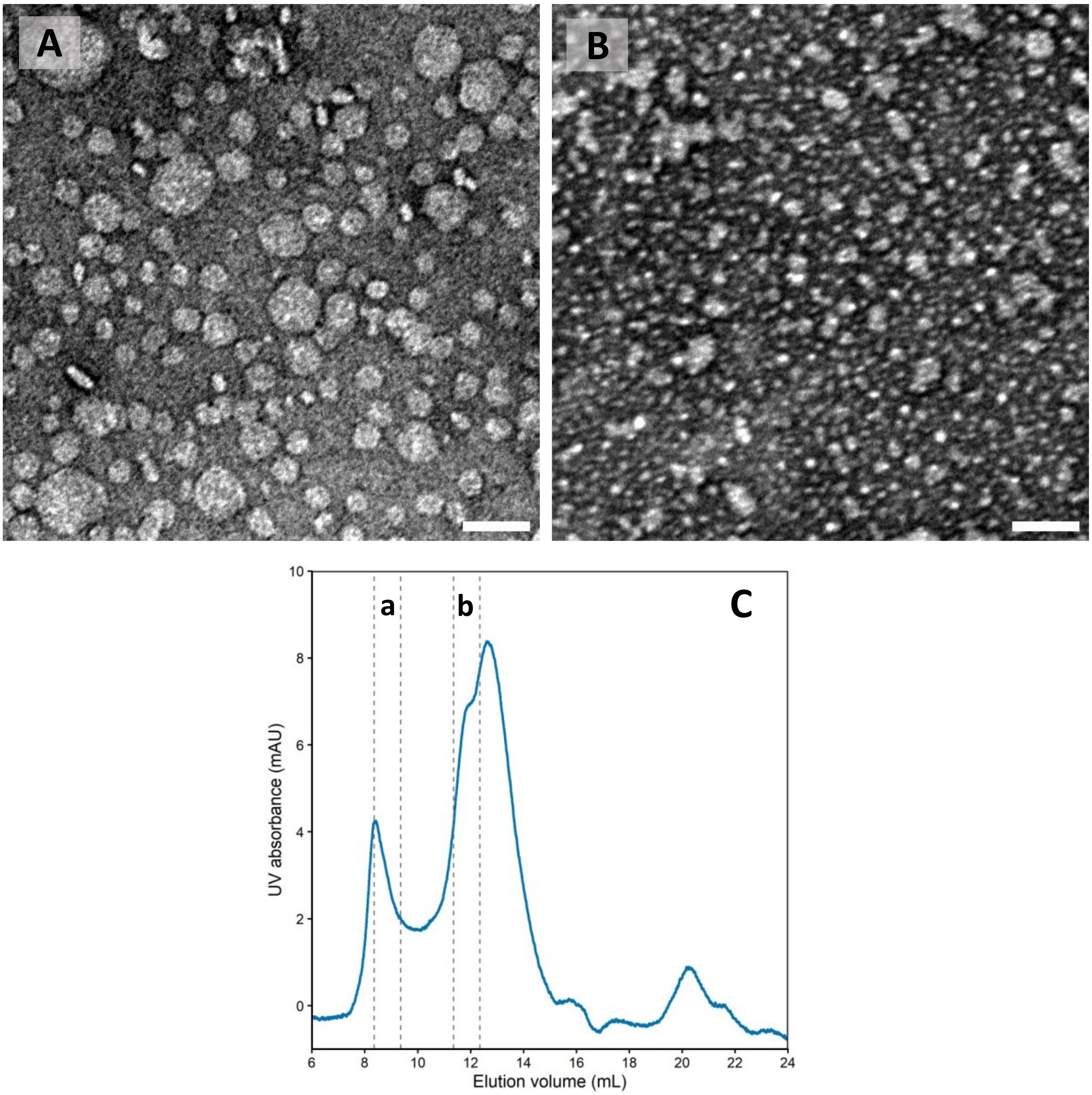


The attempt to produce ArCS1_E22TM:MSP1D1:PL nanodiscs, **(A)** representative negative-stained micrograph of fraction a, **(B)** representative negative-stained micrograph of fraction b. Micrograph scale bars, 50 nm. **(C)** SEC chromatogram of ArCS1_E22TM:MSP1D1:PL assemblies, imaged fractions are marked.

### Supplementary information 02

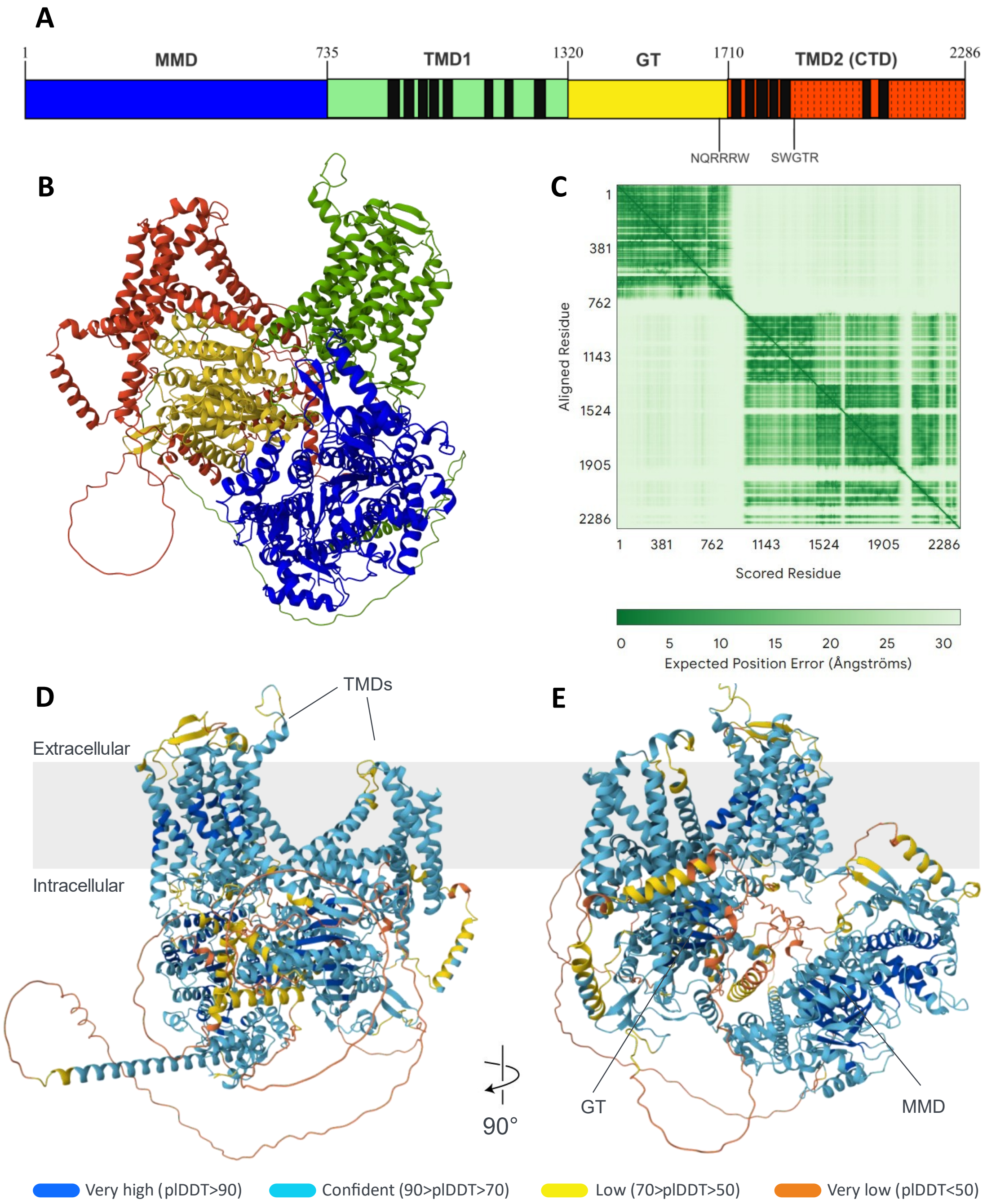


Structure of full-length ArCS1, **(A)** Domain map based on UniProt (accession number: Q288C6 ), major domains and motifs are labeled. **(B)** Cartoon representation of ArCS1 predicted by Alphafold3, colored based on the domain map. **(C)** The PAE heatmap of predicted structure. The number of residues corresponds to the amino acid sequence of the protein. **(D)** Predicted structure relative to the lipid membrane, pTM = 0.51, **(E)** with 90° rotation. The pLDDT is shown as color outputs in the structures' representations.

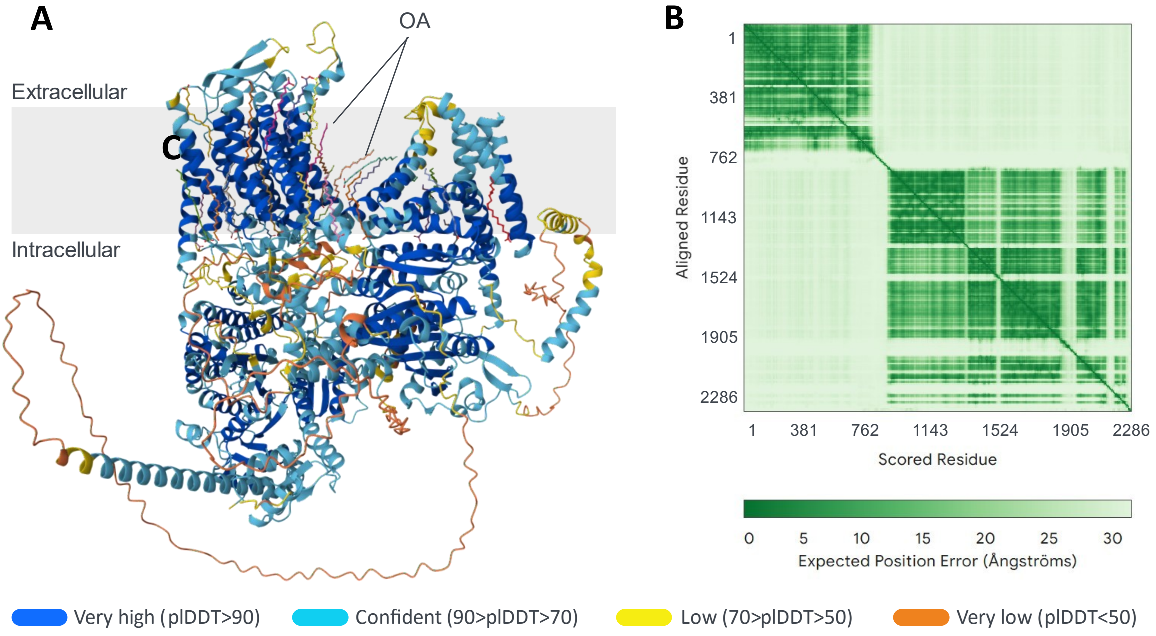


Full-length ArCS1 modeled with oleic acid (OA), **(A)** Cartoon representation relative to lipid membrane, pTM = 0.58. The pLDDT is shown as color outputs in the structures' representations. **(B)** The PAE heatmap of predicted structure. The number of residues corresponds to the amino acid sequence of the protein.
